# Immunological and clinical markers of post-acute sequelae of COVID-19: Insights from mild and severe cases six months post-infection

**DOI:** 10.1101/2025.02.18.638851

**Authors:** William Mouton, Sophia Djebali, Marine Villard, Omran Allatif, Cécile Chauvel, Sarah Benezech, Philippe Vanhems, Jacqueline Marvel, Thierry Walzer, Sophie Trouillet-Assant

## Abstract

Post-acute sequelae of COVID-19 (PASC) is a complex and multifaceted clinical challenge requiring to emphasize its underlying pathophysiological mechanisms.

This study assessed hundreds of virological, serological, immunological, and tissue damage biomarkers in two patient cohorts who experienced mild (n=270) or severe (n=188) COVID-19, 6 to 9 months post-initial infection, and in which 40% and 57.4% of patients, respectively, developed PASC.

Blood analysis showed that mains differences observed in humoral, viral, and biological biomarkers were associated with the initial COVID-19 severity, rather than being specifically linked to PASC.

However, patients with PASC displayed altered CD4^+^ and CD8^+^ memory T-cell subsets, with higher cytokine-secreting cells and increased terminally differentiated CD45RA^+^ effector memory T cells (TEMRA). Elevated SARS-CoV-2-specific T cells responsive to nucleocapsid/membrane proteins with a TEMRA phenotype were also observed. A random forest model identified these features and initial symptom duration as top variables discriminating PASC, achieving over 80% classification accuracy.

**Highlights:**

- Nine months post-initial SARS-CoV-2 infection, over 40% of patients developed PASC
- Regardless of PASC, the initial disease form influenced the main immune differences
- PASC led to altered memory T-cell subsets with elevated TEMRA phenotype
- PASC presence is associated with disease form, initial symptom count and duration

## INTRODUCTION

A significant subset of individuals who suffered from severe acute respiratory syndrome coronavirus 2 (SARS-CoV-2) infection and its associated disease, coronavirus disease 2019 (COVID-19), reports persistent symptoms well beyond the acute phase of the initial infection.^1,2^ This phenomenon, evidenced early in the pandemic, is often defined as post-acute sequelae of COVID-19 (PASC) or long COVID. In 2022, *Santé Publique France* reported a 4% prevalence of PASC in the general French adult population. In addition, a higher incidence was found among women, job seekers, and hospitalized individuals; 1.2% of patients with PASC reported a significant impact on their daily activities.^3^

PASC is characterized by a wide range of persistent symptoms and complications most frequently including respiratory issues such as dyspnea, somatic disturbances such as asthenia (fatigue or weakness), and neurological or cognitive impairments such as anosmia (loss of smell) and ageusia (loss of taste).^4,5^ Of note, manifestations can greatly vary and many other symptoms have been reported.^6–9^ In addition, PASC affects individuals across diverse demographics and health backgrounds. Altogether, these features make it a complex and multifaceted clinical challenge for healthcare systems worldwide.^10–13^

To date, various mechanisms including virus persistence, viral reactivations, persistent inflammation, neurological damage as well as immune alterations have been suggested as involved in PASC.^14–16^ Therefore, the objectives of the present study were to better understand the pathophysiological mechanisms underlying PASC and to identify biomarkers associated with PASC.

To this aim, we included two cohorts of patients who underwent mild or severe COVID-19 during the first wave of the pandemic and meticulously collected their self-reported persistent symptoms, between 6 and 9 months post-initial infection, to construct a comprehensive dataset. Statistical analyses were conducted to evaluate associations between PASC and various virological, serological, and immunological parameters measured in blood samples of patients collected from 6 to 9 months following the initial infection. Additionally, random forest classification models were used to identify key predictors of PASC within the dataset.

## MATERIAL AND METHODS

### Study populations

For the Covid-Ser cohort (patients with mild COVID-19), the clinical data were recorded by a trained Clinical Research Associate using the Clinsight software (version _ Csonline 7.5.720.1). Blood samples were processed and stored at the Centre de Ressource Biologique Neurobiotec, 69500 Bron, France. A total of 270 healthcare workers (HCWs) SARS-CoV-2 convalescent were included in a prospective longitudinal cohort study conducted in the *Hospices Civils de Lyon* (HCL, Lyon, France). Blood sampling was performed six months after infection. The Wantai SARS-CoV-2 total antibody assay, detecting anti-RBD antibodies, was used to confirm infection. Infected patients were considered to have a mild form of COVID-19 when they were symptomatic but did not require hospitalization. Written informed consent was obtained from all participants; ethics approval was obtained from the national review board for biomedical research in April 2020 (Comité de Protection des Personnes Sud Méditerranée I, Marseille, France; ID RCB 2020-A00932-37), and the study was registered on ClinicalTrials.gov (NCT04341142).

The ImmunoNosocor cohort is an extension of the NOSOCOR project registered in ClinicalTrials.gov (NCT04637867). Patients were considered to have a severe form of COVID-19 when their hospital stay was longer than 24H. The institutional review board Ile de France V approved the study on March 8, 2020 (No. 20.02.27.69817 Cat 3). Among the 189 patients hospitalized for severe COVID-19 in the HCL during the first wave of the pandemic and included in the NOSOCOR project, 188 patients were included in this study as blood sampling was not available for one patient.

Patients who agreed to participate in the study were invited for an ambulatory follow-up 6 to 9 months after onset of their initial COVID-19. During the visit, blood samples were collected and the study physician interviewed patients about the presence of PASC, defined as the persistence of at least one clinical symptom since the initial infection (e.g., dyspnea, asthenia, anosmia, agueusia, headache, and cough). A dataset of all individual measurements and clinical parameters from both cohorts was built.

### SARS-CoV-2-RBD-specific IgG antibodies

Serum levels of SARS-CoV-2-RBD-specific IgG were measured using bioMérieux Vidas® SARS-CoV-2 IgG II (9COG) diagnosis kits (bioMérieux, Marcy l’Etoile, France, #424114), according to the manufacturer’s recommendation. To standardize these assays according to the first WHO international standard, the concentrations were expressed as binding antibody units (BAU)/mL using the conversion factor provided by the manufacturer. Positive anti-RBD IgG cut-off value was provided by the manufacturer and defined at 20.33 BAU/mL.

### Anti-IFN-α2 autoantibodies

The presence of anti-IFN-α2 autoantibodies was assessed in the plasma using a commercially available kit (Thermo Fisher; Carlsbad, CA, USA, Catalog # BMS217). The positive cut-off value for autoantibodies detection was defined at 34 ng/mL, as previously described^35^. The neutralization capacity of antibodies against IFN-α2 was determined as previously described^58^. Briefly, HEK-293T cells were transfected with a plasmid encoding the firefly luciferase under the control of human Interferon-sensitive response element (ISRE) promoters. Cells were then cultured in Dulbecco’s modified Eagle’s medium (DMEM; Thermo Fisher; Carlsbad, CA, USA,) supplemented with 10% healthy control or patient’s serum/plasma, and were either left unstimulated or were stimulated with IFN-α2 (10 ng/mL) for 16 h at 37°C. Finally, luciferase level was measured using the Dual-Glo® reagent, according to manufacturer’s instructions (Promega, Madison, WI, USA).

### Circulating viral DNAemia and antigenemia

Viral DNA was extracted from 200 μL of plasma samples using the NucliSENS easyMag automated nucleic acid extractor (bioMérieux) following manufacturer’s instructions. The viral load was evaluated using quantitative real-time polymerase chain reaction (qPCR) R-Gene® assays (bioMérieux) for the following viruses: adenovirus (HAdV, Ref#69-010B), CMV (Ref#69-003B), EBV (Ref#69-002B), and TTV (Ref#69-030B), with limits of detection of 550, 290, 182, and 250 DNA copies/mL, respectively.

The concentration of SARS-CoV-2 nucleocapsid antigens was measured in plasma samples using previously developed and optimized ultra-sensitive single molecule array (SIMOA) assays on the HD-X Analyzer (Quanterix Corporation, Billerica, MA, USA), according to the protocol published by Swank *et al*^23^.

### Plasmatic protein quantification

The concentrations of plasmatic IL-6, IL-12, IL-8, monokine induced by gamma interferon (MIG/CXCL9), interferon gamma-induced protein 10 (IP-10/CXCL10), interferon gamma (IFNɣ), soluble urokinase plasminogen activator receptor (suPAR), and Vascular endothelial growth factor receptor 2 (VEGFR2) were quantified using Simple/Multi-plex cartridges (ProteinSimple, San Jose, CA, USA) using the ELLA nanofluidic system (Biotechne, Minneapolis, MI, USA), according to manufacturers’ instructions. The concentrations of plasmatic neurofilament light (Nf-L) and glial fibrillary acidic protein (GFAP) (pg/mL) were measured using SIMOA using a commercial kit (Human Neurology 2-Plex assay [N2PB], Quanterix™, Lexington, MA, USA). The assay was based on a 2-step protocol and an HD-1 Analyzer (Quanterix™). The concentrations of plasmatic IL-29/IFN-λ_1_ and IFN-β_1_ were measured using a two-plex assay on the U-PLEX Assay Platform (MesoScale Discovery (MSD), Rockville, MD, USA). The concentration of plasmatic cortisol was measured using the VIDAS® Cortisol S assay (bioMérieux).

### Spectral Flow Cytometry immunophenotyping

#### Antibodies staining

Frozen PBMCs were thawed and plated at 1–2 × 10^6^ cells per well in a 96-well U-bottom plate. Cells were then resuspended and stained with Live/Dead Fixable (ThermoFisher) for 20 min at 4 °C before staining for 45 min at room temperature (RT) with antibodies and final Fluorescence-activated cell sorting (FACS) acquisition. Antibody clones and suppliers are described in Key resources table above.

#### T cell stimulation and staining

The assay was conducted as previously described^59^. Overnight-rested PBMCs were stimulated or not with SARS-CoV-2 overlapping peptide pools against SARS-CoV-2 spike protein (PepTivator, Miltenyi Biotec, Paris, France) at a final concentration of 1 μg/mL for 1 h in the presence of 1 μg/mL monoclonal antibodies CD28 and CD49d, and then for an additional 5 h with GolgiPlug and GolgiStop (BD Biosciences). Cells were then resuspended and stained with Live/Dead Fixable (ThermoFisher) for 20 min at 4 °C before staining for 45 min at RT with antibodies against surface markers. Cells were then fixed and permeabilized using the Foxp3 kit (ThermoFisher) and stained for intracellular antigens. Single-cell fluorescence was analyzed by spectral flow cytometry with the Cytek Aurora 5L (Cytek, Fremont, CA, USA).

### Analysis of high dimensional flow cytometry datas

#### OMIQ

All Flow Cytometry Standard files (FCS) files were uploaded to the platform OMIQ from Dotmatics (www.omiq.ai, www.dotmatics.com). Manual gating was used to exclude debris, doublets, and dead cells. Then, a subsampling of 50 000 CD45^+^ alive leucocytes, or 20 000 CD4^+^, or 20 000 CD8^+^ per sample was performed. Next, we used a workflow including dimension reduction using UMAP, FlowSOM clustering analysis. For the main subset panel, FlowSOM settings were as follows: all files used, clustering features: all features used except CD45 and Live dead features; umap_1, umap_2, 25 clusters obtained with xdim = 15 and ydim = 15, rlen = 10, Distance Metric = euclidian, consensus elbow metaclustering, Random Seed =7994. For the T cell panel, FlowSOM settings were as follows: all files used, clustering features: CD45RA, CCR7, CD95, CXCR3, CD27, CD127, CD28, CCR6, CXCR5; umap_1, umap_2, 17 clusters (CD4^+^) and 7 clusters (CD8^+^) were obtained with xdim = 10 and ydim = 10, rlen = 10, Distance Metric = euclidian, consensus elbow metaclustering, Random Seed =1163 and 3281.

#### Diffcyt

Flow cytometry analysis was performed using the R package Diffcyt, that used edgeR, limma, and voom methods as part of the workflow^60^. Briefly, the FCS files were read as a Flowset. Arcsinh transformation was used with a 150cofactor. FlowSOM was used for cell clustering. For differential abundance analysis, we used diffcyt-DA-edgeR method between patients with and without PASC, and diffcyt-DS-limma for differential expression analysis^61^. We used a False discovery Rate (FDR) cut-off of <0.01 to identify differentially expressed cell types and surface markers.

### Statistical Analysis

Quantitative data were expressed as mean (range) or median [IQR] when appropriate. Normality testing was performed using the Shapiro-Wilk normality test. According to the sample distributions, parametric or non-parametric tests were used for comparison between groups comprising ANOVA (Gaussian data), or Kruskall-Wallis, or Mann-Whitney tests, when appropriate (non-Gaussian data). Correlation analyses were performed using parametric (Pearson) or non-parametric (Spearman) correlation coefficients. Adjustment for multiple testing was performed using Benjamini-Hochberg or Bonferoni correction, when appropriate. A p-value, either raw or adjusted, was considered significant when < 0.05. Plots were performed using the GraphPad Prism® software (version 8; GraphPad software, La Jolla, CA, USA) and statistical analyses were conducted using R (version 4.2.2.).

#### Machine learning analysis

Standard inferential statistical modelling was not appropriate due to the high dimensionality of potential predictors for PASC and non-PASC disease and the limited sample size of 87 observations (patients equally distributed between the two categories: Long and Non-long disease). To address this limitation, we prioritized a non-parametric method of machine learning with variable selection to identify the most impactful predictors, based on their predictive performance.

The Random Forest algorithm was used to assess the influence of all available clinical and immunological variables in discriminating PASC from non-PASC disease. The Mean Decrease in Impurity (Gini Importance) criterion was used to quantify variable importance. Variables with the highest importance scores were subsequently incorporated into a logistic regression model to identify those with statistically significant roles in predicting the binomial outcome (PASC or non-PASC).

Variable importance scores can vary depending on the randomly chosen seed values. To obtain more robust and reliable estimates of variable importance and to mitigate the influence of outlier scores, we implemented a home-made approach. This involved repeating the algorithm 1000 times, randomly selecting 500 realizations from the 1000 iterations, and aggregating the importance scores by calculating the mean of these 500 results. This process yielded stable classification performance when repeated.

## RESULTS

### Clinical characteristics of the study population during initial SARS-CoV-2 infection

The study included 458 individuals, from two cohorts of convalescent patients who experienced severe (ImmunoNosocor^17^, n=188) or mild COVID-19 (Covid-Ser^18^, n=270). All initial infections occurred during the first wave of the pandemic, between March 1^st^, and April 30^th^, 2020, during the circulation of the D614G Wuhan strain. At initial infection we observed notable clinical differences between patients who experienced severe COVID-19 and those with mild disease (Table 1). As compared to patients with mild COVID-19, there was significantly more males among patients with severe disease (59.6% *vs.* 14.6%, *p<0.001*), they were also older (median age 66.0 years [interquartile range IQR 54.3–74.0] *vs.* 40.0 years [IQR 31.0–50.0], *p<0.001*), and had a higher body mass index (BMI; 26.7 [IQR 24.3–30.4] *vs.* 23.7 [IQR 21.0–26.6], *p<0.001*). As expected, patients suffering from a severe COVID-19 also had a higher prevalence of comorbidities (79.8% *vs.* 35.9%, *p<0.001*), including cardiovascular disorders and heart failure (*p<0.001)*.

**TABLE 1.** Demographic and clinical characteristics of patients with mild or severe acute SARV-CoV-2 infection, Lyon University Hospitals, France.

| Initial form of COVID-19<br>Status, n (%) | Severe<br>All, 188 | Mild<br>All, 270 | *p.value*<sup>a</sup> | Severe<br>w/ PASC, 108 | w/o PASC 80 | *p.value*<sup>a</sup> | Mild<br>w/ PASC, 108 | w/o PASC, 162 | *p.value*<sup>a</sup> |
| --- | --- | --- | --- | --- | --- | --- | --- | --- | --- |
| **Demographical data** |  |  |  |  |  |  |  |  |  |
| Male, n (%) | 112 (59.6) | 40 (14.8) | **<0.001** | 62 (57.4) | 50 (62.5) | 0.48 | 11 (10.2) | 29 (17.9) | 0.116 |
| Age at blood sampling, median [IQR] | 66 [54-74] | 40.0 [31-50] | **<0.001** | 62.6 [25-93] | 64.4 [28-84] | 0.23 | 41.5 [33-50] | 38.0 [30-50] | 0.396 |
| Body Mass Index, median [IQR] | 26.7 [24-30] | 23.7 [21-27] | **<0.001** | 28.3 [25-31] | 25.9 [24-28] | **0.011** | 23.8 [21-27] | 23.5 [21-26] | 0.400 |
| Currently smoker, n (%) | 4 (2.1) | 29 (10.7) | **<0.001** | 1 (0.93) | 3 (3.8) | 0.09 | 16 (14.8) | 13 (8.0) | 0.281 |
| Daily alcohol consumption, daily n (%) | 6 (3.2) | 4 (1.5) | 0.330 | 3 (2.8) | 3 (3.8) | 0.74 | 3 (2.8) | 1 (0.62) | 0.486 |
| **Comorbidities** |  |  |  |  |  |  |  |  |  |
| Presence, n (%) | 150 (79.8) | 97 (35.9%) | **<0.001** | 84 (77.8) | 66 (82.5) | 0.53 | 43 (39.8) | 54 (33.3) | 0.338 |
| Cardiovascular disorders, n (%) | 84 (56.0) | 9 (9.3) | **<0.001** | 49 (45.4) | 35 (43.8) | 0.83 | 4 (3.7) | 5 (3.1) | 1.000 |
| Chronic pulmonary disorders, n (%) | 12 (8.0) | 17 (17.5) | **0.039** | 7 (6.5) | 5 (6.3) | 0.95 | 8 (7.4) | 9 (5.6) | 0.720 |
| Neurological disorders, n (%) | 9 (6.0) | 8 (3.0) | 0.672 | 3 (2.8) | 6 (7.5) | 0.13 | 4 (3.7) | 4 (2.5) | 0.717 |
| Hypertension, n (%) | 67 (44.7) | 18 (18.6) | **<0.001** | 38 (35.2) | 29 (36.3) | 0.88 | 6 (5.6) | 12 (7.4) | 0.727 |
| Heart failure, n (%) | 24 (16.0) | 1 (1.0) | **<0.001** | 12 (11.1) | 12 (15) | 0.43 | 1 (0.9) | 0 (0.0) | 0.400 |
| Diabetes, n (%) | 36 (24.0) | 7 (7.2) | **0.001** | 18 (16.7) | 18 (22.5) | 0.32 | 3 (2.8) | 4 (2.5) | 1.000 |
| Cancer, n (%) | 21 (14.0) | 6 (6.2) | 0.087 | 13 (12) | 8 (10) | 0.66 | 2 (1.9) | 4 (2.5) | 1.000 |
| Immune deficiency, n (%) | 10 (6.7) | 1 (1.0) | 0.054 | 5 (4.6) | 5 (6.3) | 0.62 | 0 (0.0) | 1 (0.6) | 1.000 |
| Liver disease, n (%) | 16 (10.7) | 1 (1.0) | **0.008** | 11 (10.2) | 5 (6.3) | 0.33 | 0 (0.0) | 1 (0.6) | 1.000 |
| Kidney disease, n (%) | 18 (12.0) | 4 (4.1) | 0.097 | 9 (8.3) | 9 (11.3) | 0.50 | 3 (2.8) | 1 (0.6) | 0.305 |
| Hypothyroidy, n (%) | 17 (11.3) | 15 (15.5) | 0.580 | 12 (11.1) | 5 (6.3) | 0.25 | 6 (5.6) | 9 (5.6) | 1.000 |
| Rheumatic disease, n (%) | 20 (13.3) | 10 (10.3) | 0.754 | 13 (11.9) | 7 (8.8) | 0.48 | 5 (4.6) | 5 (3.1) | 0.527 |
| **Clinical data during initial infection** |  |  |  |  |  |  |  |  |  |
| Symptoms number, median [IQR] | 5 [4-7] | 8 [6-10] | **<0.001** | 6 [4-7] | 5 [3-6] | 0.173 | 9 [7-12] | 7 [5-9] | **<0.001** |
| Symptoms duration (days), median [IQR]<sup>b</sup> | 19 [13-28] | 11 [7-26] | **<0.001** | 21 [14-29] | 17 [12-22] | **0.008** | 29 [11-80] | 10 [7-16] | **<0.001** |
| Dyspnea, n (%) | 123 (65.4) | 128 (47.4) | **<0.001** | 77 (71.3) | 46 (57.5) | 0.088 | 64 (59.3) | 64 (39.5) | 0.002 |
| Asthenia, n (%) | 138 (73.4) | 233 (86.3) | **0.002** | 80 (74.1) | 58 (72.5) | 1.000 | 102 (94.4) | 131 (80.9) | 0.003 |
| Anosmia, n (%) | 26 (13.8) | 150 (55.6) | **<0.001** | 19 (17.6) | 7 (8.8) | 0.136 | 71 (65.7) | 79 (48.8) | 0.009 |
| Agueusia, n (%) | 25 (13.3) | 133 (49.3) | **<0.001** | 14 (13.0) | 11 (13.8) | *1.000* | 61 (56.5) | 72 (44.4) | *0.070* |
| Headaches, n (%) | 38 (20.2) | 188 (69.6) | **<0.001** | 22 (20.4) | 16 (20.0) | *1.000* | 87 (80.6) | 101 (62.3) | *0.002* |
| Cough, n (%) | 149 (79.3) | 147 (54.4) | **<0.001** | 87 (80.6) | 62 (77.5) | *0.870* | 68 (63.0) | 79 (48.8) | *0.030* |
| ICU admission, n (%) | 44 (23.4) | NA | *NA* | 32 (29.6) | 11 (13.8) | **0.03** | NA | NA | *NA* |
| ICU length of stay (days), median [IQR] | 27 (13-45) | NA | *NA* | 23.4 [1-184] | 14.9 [1-128] | **0.004** | NA | NA | *NA* |

<sup>a</sup>Comparisons were performed using Fisher's exact test or using the non-parametric Mann-Whitney U test based on the sample distributions. \*, p < 0.05; \*\*, p < 0.01; \*\*\*, p < 0.001. P.values are italicized and bolded when they reach a significant value. <sup>b</sup>Missing value for 25/188 patients of severe cohort and 159/270 patients of mild cohort. *Abbreviation: ICU, intensive care unit; IQR, interquartile range; w/, with; w/o, without.*

### Clinical characteristics of patients suffering from PASC

Among the 458 patients included, 216 patients (108 from each cohort) self-reported a lack of return to their pre-COVID-19 health status with at least one persistent symptom between 6- and 9-months post-infection (median [IQR] 189 days [183–198] for mild COVID-19 and 290 days [267–310] for severe COVID-19; Table 1); these patients were therefore classified as patients with PASC.

PASC was less prevalent following mild COVID-19 compared to severe COVID-19 (40.0% *vs.* 57.4%, *p<0.001*). Among persistent symptoms, anosmia was significantly more frequent in mild-PASC patients than in severe-PASC patients (33.3% vs. 11.1%; *p<0.001*). Other symptoms, such as dyspnea (37.0% vs. 35.5%) and asthenia (33.3% vs. 34.3%), were reported at similar rates in both cohorts (Figure 1A).

**Figure 1:**
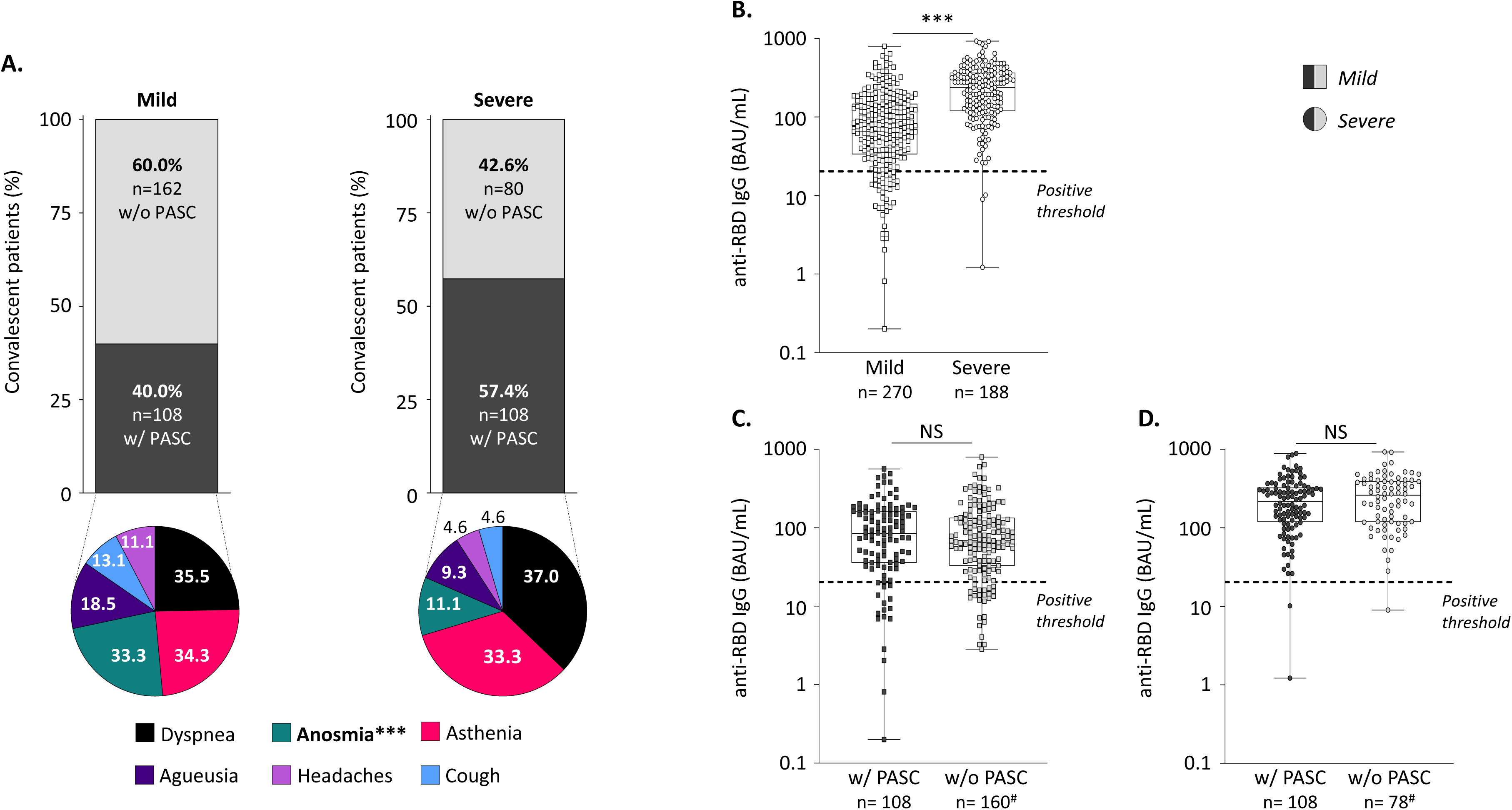
Persistent symptoms and anti-RBD IgG levels in patients with or without PASC. Included patients were classified according to their initial form of COVID-19, whether severe or mild, and the persistence of symptoms (with or without PASC), as indicated. (A) Proportions of PASC in each cohort, and of persistent symptoms self-reported during the interview. Comparisons were performed using Fisher’s exact test. (B) Serum anti-RBD IgG levels according to the severity of initial COVID-19 episode. Serum anti-RBD IgG levels according to PASC status in patients with initial mild (C) and severe (D) COVID-19 episode. Results are expressed as BAU/mL and the positivity cut-off value applied was defined by the manufacturer at 20.33 BAU/mL. Comparisons were performed using the non-parametric Mann-Whitney U test. *, p < 0.05; **, p < 0.01; ***, p < 0.001. *Abbreviation: BAU, binding antibody unit; IgG, immunoglobulin G; RBD, receptor-binding domain.* #Data were missing for two patients in both cohorts.

As previously described,^19^ in the severe cohort, PASC was associated with a higher BMI (28.3 [25-31] vs. 25.9 [24-28], *p=0.011*), longer initial symptom duration (21 [14-29] vs. 17 [12-22] days, *p=0.008*), a greater proportion of ICU admissions (29.6% vs. 13.8%, *p=0.03*), a longer ICU length of stay (23 [1-184] vs. 15 [1-128] days, *p=0.004*), and more frequent complications during hospital stays, such as respiratory issues (31.5% vs. 13.8%, *p=0.019*) or undergoing ventilation (31.5% vs. 16.3%, *p=0.024*), which reflects the severity of the acute initial phase. We subsequently conducted laboratory investigations on blood samples collected from 6 to 9 months after the initial infection to explore the pathophysiological mechanisms underlying PASC. Of note, all patients were not vaccinated at the time of blood sampling.

### SARS-CoV-2 anti-RBD IgG antibodies

Although a direct link between specific IgG antibody levels and PASC has not been established yet, emerging evidence suggests that individuals with PASC, from 1 to 9 months after initial infection, exhibit significantly elevated levels of anti-SARS-CoV-2 IgG compared to those without persistent symptoms.^14,20–22^ Therefore, our first objective was to explore potential differences in the humoral immune response among both cohorts. Regardless of the PASC status we showed a significantly higher levels of anti-RBD IgG in patients severe COVID-19 compared to those with mild disease (237.9 [119.1-365.7] *vs.* 75.4 [33.6-147.3] BAU/mL, respectively, *p=0.0001*). We also observed a higher proportion of individuals above the IgG positive detection cut-off value among severe patients in comparison to mild patients (98.4% *vs.* 84.7% respectively, *p<0.0001*; Figure 1B). Nevertheless, when comparing patients with PASC to those without, our analysis showed no significant difference in anti-RBD IgG antibody levels; 84.8 [35.9-161.4] *vs.* 68.9 [32.9-133.7] BAU/mL, respectively, *p=0.40* for the mild cohort (Figure 1C) and 217.4 [119.4-324.9] *vs.* 260.1 [119.1-393.8] BAU/mL, respectively, *p=0.24* for the severe cohort (Figure 1D). Similar observations were made regarding the proportion of patients above the positive detection cut-off value in the mild cohort (85.2% *vs.* 84.4%, respectively, *p>0.99*) and in the severe cohort (98.1% *vs.* 98.7%, respectively, *p>0.99*). Of note, all these observations remained consistent after statistical adjustment for the time between symptom onset at the initial infection and blood sampling.

### Detection of anti-IFN-I autoantibodies and circulating viral DNAemia and antigenemia

Previous studies reported a link between PASC symptoms and plasmatic factors, including anti-interferon-α (IFNα) autoantibodies,^14,15^ SARS-CoV-2 nucleocapsid antigens,^23,24^ or various reactivated viruses.^15,25,26^ We therefore investigated these factors and observed a low prevalence of anti-IFN-α2 plasmatic autoantibodies with neutralizing capacity; among the 458 patients, they were found in only 6 severe patients (1.3%) without association with PASC. Regarding circulating SARS-CoV-2 nucleocapsid antigens, they were not detectable in any of the samples. Similarly, Cytomegalovirus (CMV) and adenovirus were not detectable, while only 10 samples were positive for Epstein-Barr virus (EBV; 2.2%), without association with PASC. Conversely, torque teno virus (TTV) load was detectable in most samples and was significantly different between cohorts; severe patients had a higher viral load than mild patients (3.05 [2.23–3.73] vs. 2.43 [1.80–3.00] Log₁₀ copies/mL, *p<0.0001*). However, while a higher proportion of TTV-positive samples was found in severe patients with PASC compared to those without (58% vs. 42%, *p=0.0068*; Table S1), no significant difference in TTV loads was observed according to the PASC status.

### Plasma cytokines and biomarkers of tissue damage

Several studies have found that individuals with PASC have elevated serum levels of pro-inflammatory cytokines, such as IFNα, Interleukin(IL)-6, IL-1β, and TNF-α, or chemokines such as CXCL10.^27–29^ Chronic inflammation may cause tissue damage, as evidenced in several studies showing the presence of brain or vascular antigens in the plasma. To address this question in our study, we first selected subgroups in each cohort in order to match patients with and without PASC for age, sex, and comorbidities. A total of 90 subjects were selected in the severe cohort (45 with PASC and 45 without PASC) and 85 in the mild cohort (45 with PASC and 40 without PASC). We then quantified the plasmatic levels of cytokines IL-12, IL-6, IL-8, IFN-β1, IFN-λ1, IFN-γ, and chemokines CXCL9 or CXCL10 in these 175 patients. No significant difference in cytokine levels was observed between patients with and without PASC (Figure 2A-H). A similar finding was obtained upon investigation of serum levels of biomarkers related to endothelial damage, (soluble urokinase plasminogen activator receptor [suPAR] and vascular endothelial growth factor receptor 2 [VEGFR2], Figure 2I-J), or neuronal damage (neurofilament light [Nf-L] and Glial fibrillary acidic protein [GFAP], Figure 2K-L). Interestingly, levels of several cytokines (IL-6, IFN-λ1, CXCL9, and CXCL10) and tissue damage biomarkers were greater in patients who experienced severe COVID-19 compared to those with a mild form, regardless of PASC status; depending on the cytokine and biomarker, levels ranged from 1.2 to 1.6 times greater, and *p-values* from 0.0011 to <0.001; Figure S1).

**Figure 2:**
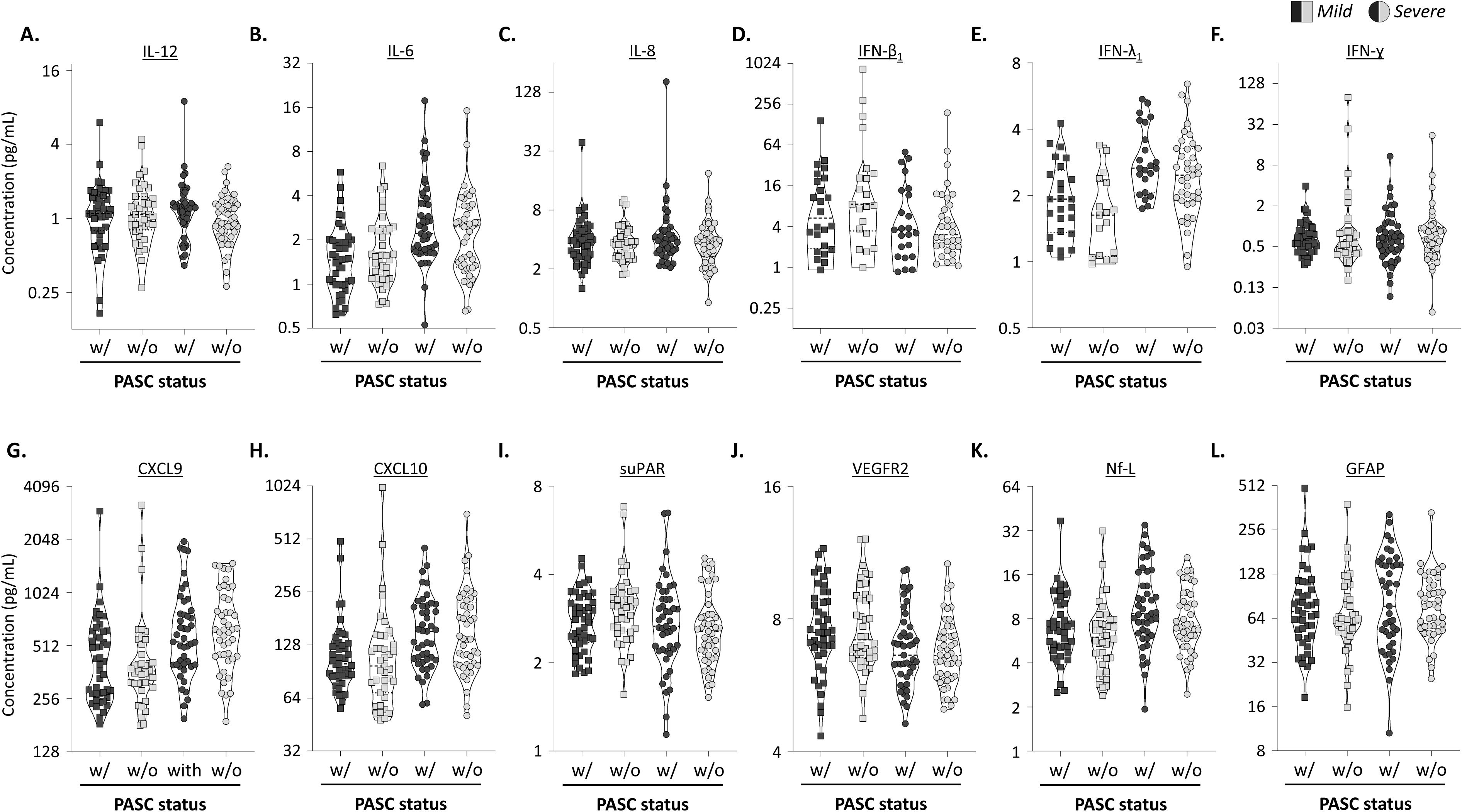
Plasmatic levels of cytokines and tissue damage biomarkers according to PASC status and severity of the initial COVID-19 episode. Plasmatic levels of IL-12 (A), IL-6 (B), IL-8 (C), IFN-β_1_ (D), IFN-λ_1_ (E), IFN-γ (F), CXCL9 (G), CXCL10 (H), suPAR (I), VEGFR2 (J), Nf-L (K), and GFAP (L). Comparisons were performed between patients grouped according to severity of initial COVID-19 and PASC status (mild COVID-19: squares, severe COVID-19: circles, with PASC: black symbols, without PASC: grey symbols). Results were expressed as pg/mL. #Levels were below the limit of detection for IFNβ_1_ in 40 of the 85 mild patients and 35 of the 90 severe patients, and for IFNλ_1_ in 43 of the 85 mild patients and 28 of the 90 severe patients, respectively. Comparisons were performed using the non-parametric Mann-Whitney U test. *, p < 0.05; **, p < 0.01; ***, p < 0.001. *Abbreviation: GFAP, Glial fibrillary acidic protein; IFN, interferon; IL, interleukin; IP, interferon protein; MIG, monokine induced by gamma interferon; Nf-L, neurofilament light; suPAR, soluble urokinase plasminogen activator receptor; VEGFR2, vascular endothelial growth factor receptor 2*.

### Level of circulating cortisol

Cortisol levels have been recently investigated in patients with PASC; Klein *et al.*^14^ found significantly lower cortisol levels in these patients, which was not confirmed Fleischer *et al.*^30^ who did not report such a difference. Among the present patients, cortisol was quantified in 160 of the available 175 samples (with enough volume), and significantly higher levels were observed in severe patients, as compared to mild patients (90.9 [73.9–124.4] vs. 74.9 [60.9–101.4] pg/mL, *p=0.0045*; Figure S2A).

However, no significant difference in cortisol levels was found between patients with and without PASC, neither in the mild (93.6 [63.4–109.2] vs. 71.2 [55.3–93.4] pg/mL, p=0.070; Figure S2B) nor in the severe cohort (90.4 [67.6–125.3] vs. 100.3 [80.4– 118.3] pg/mL, *p=0.38*; Figure S2C).

### Patients with and without PASC display differences in the phenotypes of memory CD4^+^ and CD8^+^ T cell subsets

Since the main differences observed between patients were related to the severity of the initial disease, and as a higher proportion of PASC was found in patients who had experienced severe COVID-19, we decided to focus the following analyses exclusively on this cohort. We performed immunophenotyping using peripheral blood mononuclear cells (PBMC) samples from 88 patients of the severe cohort (44 with PASC vs. 44 without PASC) obtained 6 months after the primary infection (Figure 3A). Two antibody panels were used to characterize either the phenotype of the overall PBMC population (Main subsets), or the phenotype of CD4^+^ and CD8^+^ T cells (T cell panel; Table S2) using flow cytometry. The analysis of main cell subsets identified through manual gating (Figure S3) revealed limited differences between patients according to the PASC status. Specifically, patients with PASC tended to exhibit higher counts of lymphocytes, while patients without PASC tended to exhibit higher counts of total and classical monocytes, IgD^+^CD27^+^ B cells, and CD45RA^-^/CD95^+^ CD8^+^ T cells (Figure 3B). These results highlights notable alterations in T cell counts, which were previously reported in patients with PASC,^14,15,31^ therefore, we performed unsupervised clustering of T cell data using the FlowSom algorithm to further explore T cells in the context of PASC. This analysis identified 17 distinct clusters of CD4^+^ T cells and 7 clusters of CD8^+^ T cells (Figure S4). We then leveraged the diffcyt algorithm to compare the mean expression of each marker between patients with and without PASC (Table S3). Several markers were differentially expressed between the two groups, mainly on memory CD4^+^ and CD8^+^ T cells. Among memory CD4^+^ T cells, effector memory T cells re-expressing CD45RA (TEMRA, cluster 17) and effector memory T cells (TEM, cluster 10) were observed to have a similar abundance but distinct phenotypes depending on the PASC status (Figure 4A). In patients with PASC, CD4^+^ TEMRA cells (cluster 17) showed higher expression of CD57 (*p=0.015*) and Tbet (*p=0.048*) as well as lower CXCR3 expression (*p=0.015*). Furthermore, CD4^+^ TEM cells (cluster 10) in these patients showed reduced CD27 expression (*p=0.026*; Figure 4A). Similarly, CD8^+^ TEMRA cells (cluster 5) from patients with PASC displayed higher Tbet expression (*p=0.032*) and lower CXCR3 expression (*p=0.0072)*, despite a similar abundance found among patients with and without PASC (Figure 4B). We found that slight immune alterations of CD4^+^ T-cell phenotype discriminated patients with PASC from those without, although these alterations did not represent a common feature for all patients with PASC.

**Figure 3:**
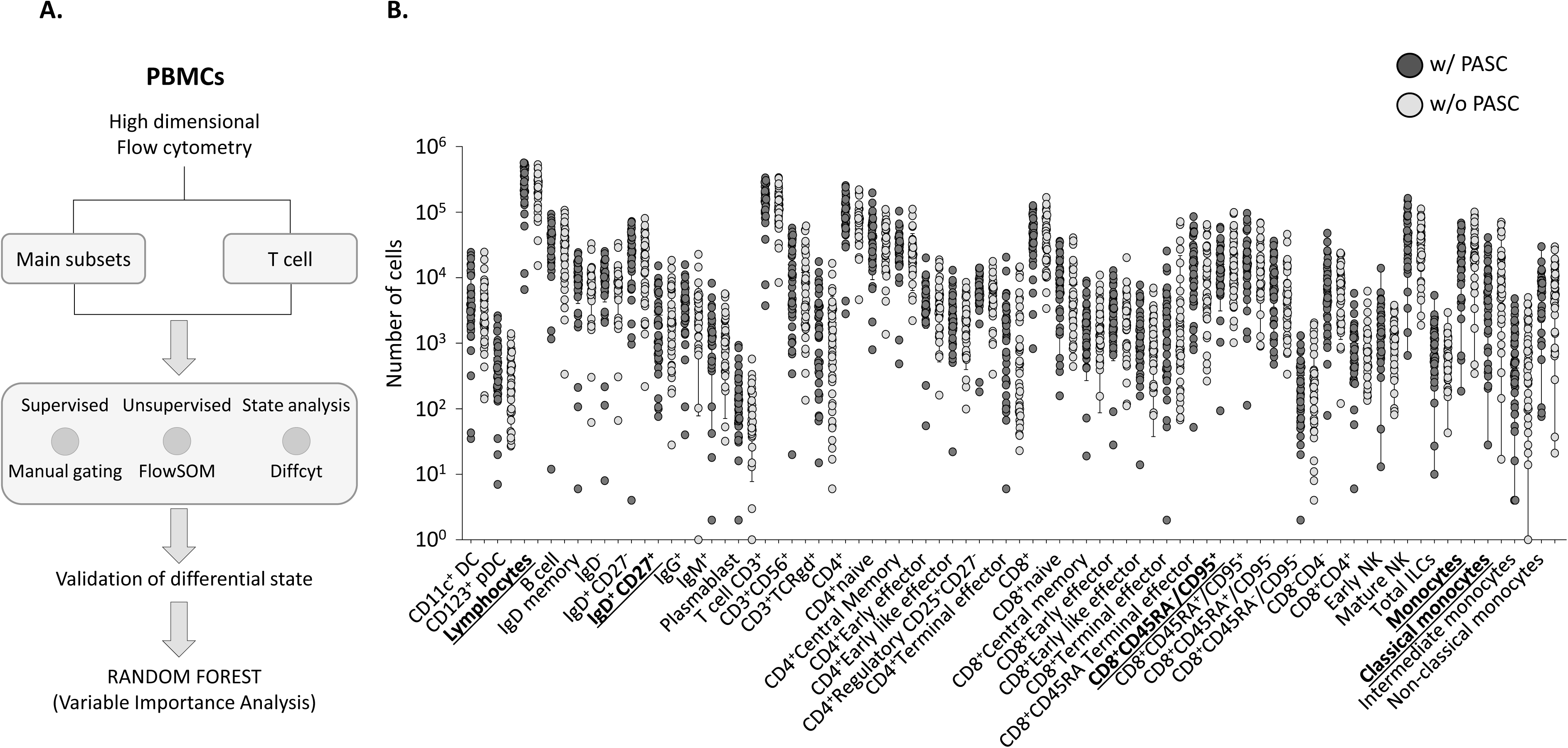
Flow cytometry analysis of immune subsets from 6 months post-infection. (A) Flowchart of the cytometry data analysis: PBMCs were labelled using high dimension flow cytometry panels allowing the analysis of main subsets within PBMCs as well as the detailed phenotype of T cells. Manual gating or FlowSOM unsupervised analysis were performed on the flow cytometry analysis platform OmiQ. The differential state of cell clusters was analyzed using the diffcyt algorithm and the R software. (B) Cellular count for the different immune populations defined by manual gating, comparing patients with and without PASC. Statistical analysis was performed using the non-parametric Mann-Whitney U test. *, p < 0.05; **, p < 0.01; ***, p < 0.001. *Abbreviation: CD, cluster of differentiation; Ig, immunoglobulins; ILC, innate lymphoid cell; NK, natual killer; PASC, post-acute sequalae of COVID-19*

**Figure 4:**
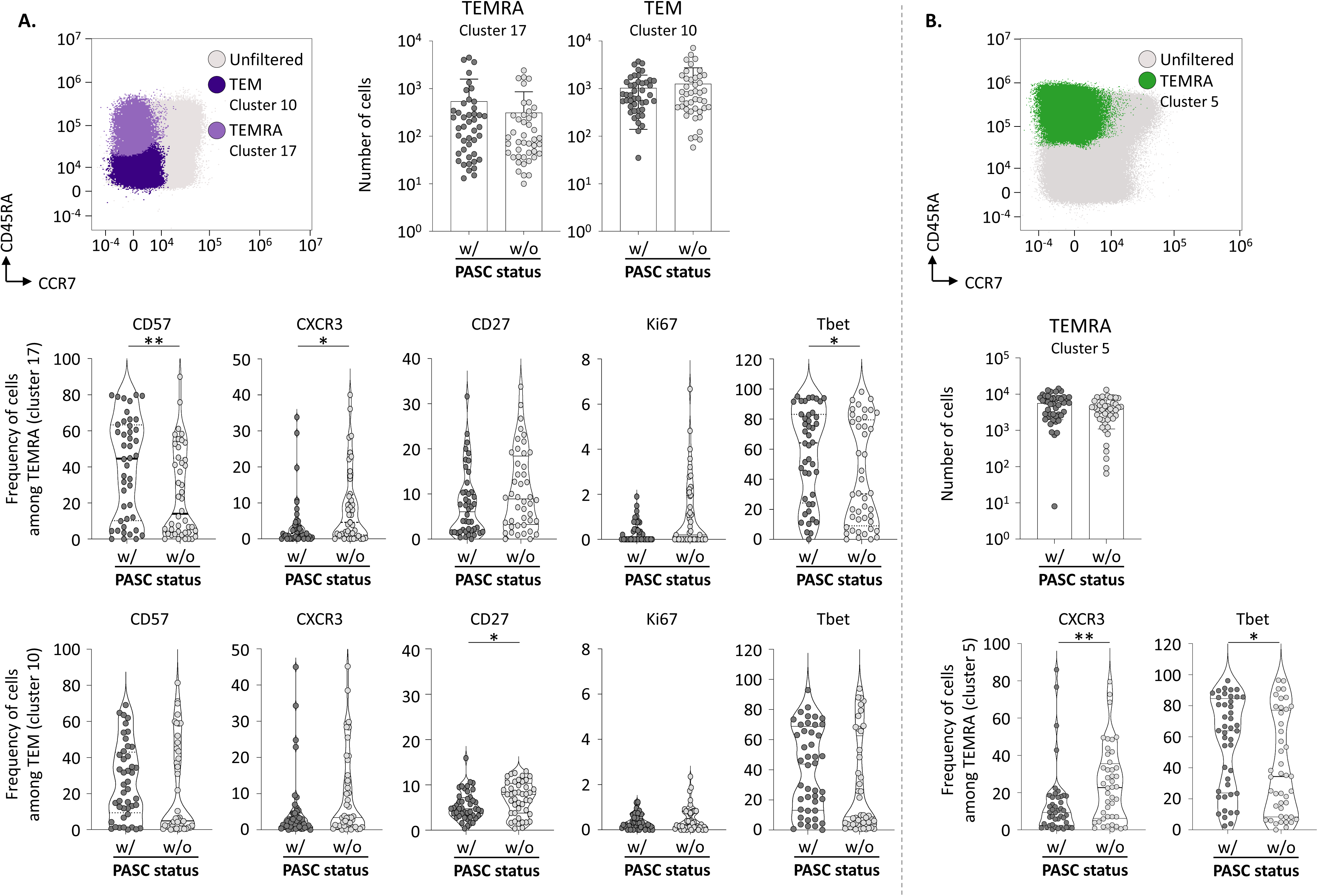
Differential T cell profiles in patients with and without PASC. PBMCs from patients with (44 samples) and without (44 samples) PASC were analyzed by spectral flow cytometry using the “T cell” antibody panel. Unsupervised clustering was performed using FlowSOM and differential expression of phenotypic markers was defined using diffcyt. Results presented in the figure highlight the most significant differences according to PASC status. Among CD4^+^ T cells (A) and CD8^+^ T cells (B) CD45RA/CCR7 expression; abundance of cells per clusters; and frequency of cells expressing the indicated markers among identified T cell clusters are shown. Data were represented as mean ± SD. Statistical analysis was performed using the non-parametric Mann-Whitney U test. *, p < 0.05; **, p < 0.01; ***, p < 0.001. *Abbreviation: CD, cluster of differentiation;* CXCR, *C-X-C chemokine receptor; PBMC, peripheral blood mononuclear cells; TEM, Effector memory cells; TEMRA, terminally differentiated effector memory*

### SARS-CoV-2 specific T cell responses are greater in patients with PASC

We next assessed T cell responses to SARS-CoV-2 proteins, specifically the spike (S) or a combination of nucleocapsid and membrane (N/M) proteins, at the single-cell level using flow cytometry. PBMCs from patients with or without PASC were stimulated, or not, with a pool of commercial 15-mer and 11-mer peptides covering the entire proteins and designed to stimulate both CD4^+^ and CD8^+^ T cells, and were then stained for intracellular IFNγ and tumor necrosis factor α (TNFα). The frequency of cytokine-secreting T cells was evaluated using the gating strategy presented in Figure S5. Regarding the unstimulated condition, we found a significantly higher proportion of cytokine-secreting T cells in patients with PASC (Figure 5A). The response to S was similar in both groups, but patients with PASC exhibited a significantly higher response to N/M-derived peptides (Figure 5A). To account for the numerous factors that could influence the frequency of cytokine-secreting T cells in response to the N/M-derived peptides, we used a multiple logistic regression model to evaluate the impact of various biological parameters, including sex, age, weight, initial symptom duration, ICU admission, ventilation, comorbidities, number of symptoms, and PASC status. The analysis revealed that the PASC status was the only parameter significantly associated with the T cell response to N/M in the cohort (*p=0.045;* Table S4).

**Figure 5:**
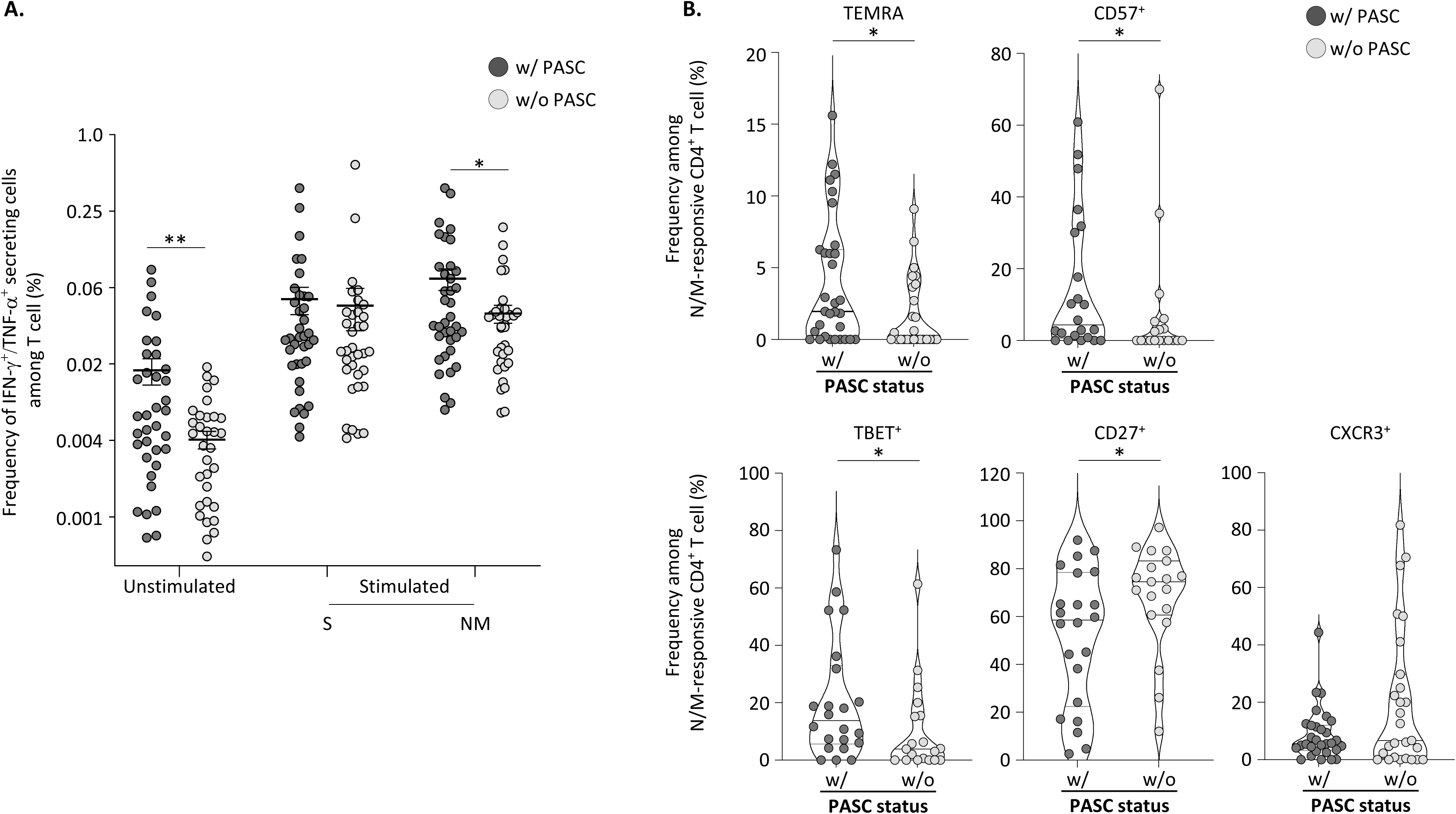
Distribution and phenotypic characteristics of SARS-CoV-2-specific CD4^+^ and CD8^+^ memory T cell responses. PBMCs from PASC and non-PASC patients were incubated, or not, with SARS-CoV-2 peptide pools from the S or N and M proteins, and T cell cytokine production was analyzed by flow cytometry. (A) Frequency of IFN-γ^+^/TNF-α^+^ secreting cells at basal state or following stimulation with S or N/M peptide pools, (B) Relative proportions of TEMRA, CD57, Tbet, CD27, and CXCR3 positive cells within N/M-responsive CD4^+^ T cells according to PASC status. To define the phenotype of SARS-CoV-2-T cells, a cut-off of 15 events was used to ensure the accuracy of the measurements. Statistical analysis was performed using the non-parametric Mann-Whitney U test. *, p < 0.05; **, p < 0.01; ***, p < 0.001. *Abbreviation: CD, cluster of differentiation;* CXCR, *C-X-C chemokine receptor; IFN, interferon; NM, nucleocapsid and membrane; PBMC, peripheral blood mononuclear cells; S, spike; TEMRA, terminally differentiated effector memory; TNF, tumor necrosis factor*.

We then compared the proportion of CD4^+^ and CD8^+^ T cells responding to N/M or S, between patients with and without PASC, using concatenated flow cytometry data. In patients with PASC, there was a trend toward a greater proportion of CD4^+^ T cells among N/M-responsive T cells, but not among S-responsive T cells (Figure S6A and S6B). Finally, we compared the phenotype of N/M-responsive CD4^+^ and CD8^+^ T cells using other surface markers available in our flow cytometry panel. We found that the N/M-responsive CD4^+^ T cells exhibited a higher frequency of TEMRA cells, and expressed higher levels of CD57 and Tbet, but lower levels of CD27 and CXCR3 (Figure 5B) in patients with PASC compared to patients without. No significant difference was observed in N/M-responsive CD8^+^ T cells when comparing both patient groups (Figure S6C).

### Classification of parameters discriminating patients with and without PASC

In order to find the variables that best discriminated patients with PASC from those without, we used a random forest algorithm as well as the 674 clinical, serological, virological, and immunological parameters monitored in this study. Figure 6A and Figure S7 illustrate the top 10 and top 50 discriminant parameters. This list included clinical (symptom duration, weight), plasmatic (GFAP, IL-6, Nf-L, and IFN-λ1) as well as immunological parameters (CD8^+^ TEMRA CXCR3, CD4^+^ TEMRA Ki67, N/M-responsive CD4^+^ TEMRA, classical monocytes). The most important parameter identified was the symptom duration, followed by several immune parameters associated with T cells. However, none of these parameters alone was sufficient to fully differentiate patients with PASC from those without. We then assessed the capacity of the top 10 parameters to correctly discriminate altogether patients with and without PACS, using a confusion matrix. As shown in Figure 6B these 10 parameters had a classification performance greater than 80% for both group of patients, supporting a role for T cells in PASC.

**Figure 6:**
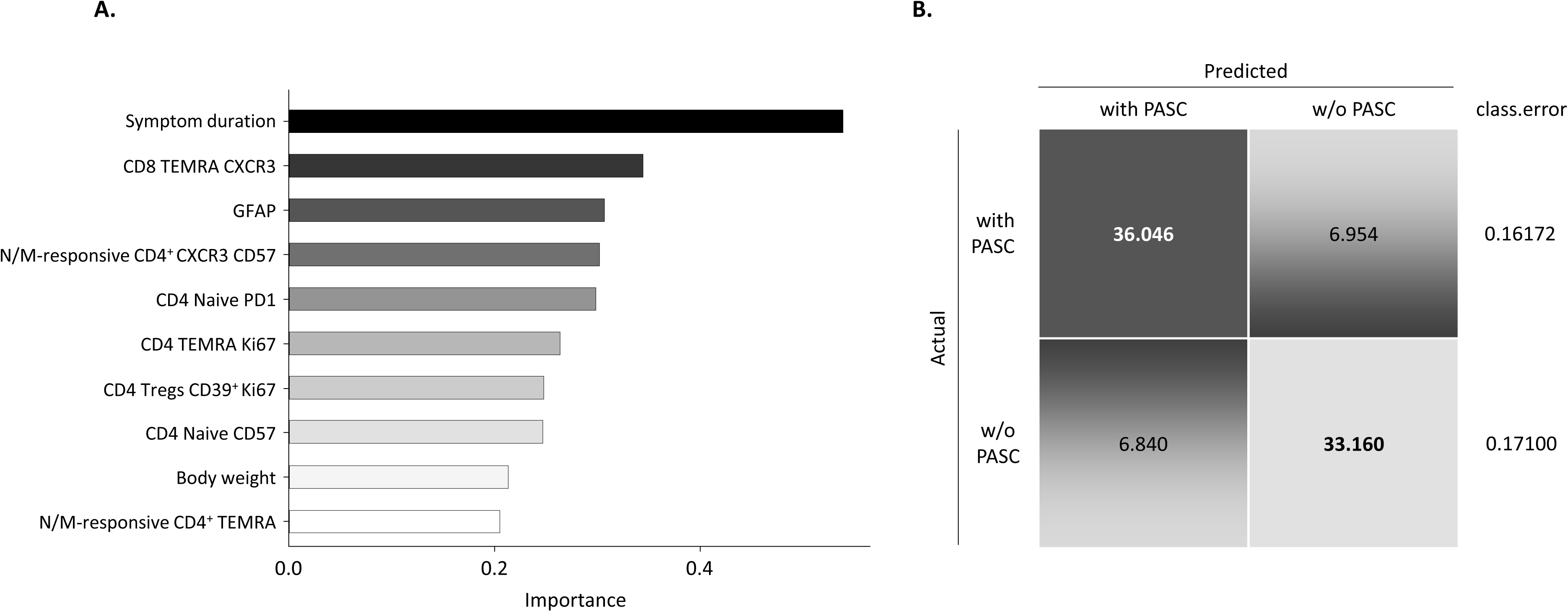
Evaluation of clinical and biological parameters to discriminate patients according to PASC status. (A) A random forest approach was applied using clinical and biological parameters assessed in this study to identify the factors that best discriminate patients with PASC from those without. The top 10 parameters are shown. Predictor variables were ranked based on the Gini Importance index, calculated using the Random Forest algorithm within the R statistical environment. Higher scores indicate a stronger contribution to predict the dichotomous outcome (PASC vs. non-PASC). (B) The confusion matrix showing the classification performance of these 10 parameters. The matrix confronts the conditions predicted by the algorithm to the true conditions. Each row of the matrix represents the patient status (with or without PASC), while each column represents the predicted status. Results were expressed as a percentage (%). The mean error rate on the 1000 iterations of the random forest is given for each group of patients. *Abbreviation: Ag, antigen; CD, cluster of differentiation;* CXCR, *C-X-C chemokine receptor; GFAP, Glial fibrillary acidic protein; Sp, specific TEM, Effector memory cells; TEMRA, terminally differentiated effector memory*

## DISCUSSION

The present study offers a comprehensive analysis of two cohorts of patients with mild or severe COVID-19, monitored at diagnosis and assessed for PASC from 6 months post-infection. Our analysis highlighted differences in clinical and immunological parameters such as SARS-CoV-2 anti-RBD IgG, cortisol, or cytokines (IL-6, IFN-λ1, CXCL-10, CXCL-9) levels between mild and severe cohorts but not according to the PASC status. However, an in-depth analysis of immune parameters in severe patients with and without PASC, highlighted that total and N/M-responsive memory T cells from patients with PASC harbored a more terminally differentiated phenotype. The association of these cellular phenotypes with the development of PASC was supported by a random forest classifier that identified the most important parameters associated with PASC.

While the prevalence of PASC reported in studies conducted at least 12 weeks after infection ranges from 0% to 93%,^32^ nearly half of the individuals herein reported PASC. Consistent with previous observations,^5,9,27,33^ a higher proportion of patients with PASC was found following a severe initial COVID-19, although the distribution of the various persistent symptoms reported was similar in both cohorts.

Our analysis found that patients with severe COVID-19 had significantly higher levels of SARS-CoV-2 anti-RBD IgG compared to those with mild COVID-19. However, in both cohorts no significant difference in anti-RBD IgG levels was found according to the PASC status. This highlights that the humoral response fluctuation is predominantly influenced by the initial COVID-19 form but may not correlate with the presence of persistent symptoms, as already suggested.^15,28,34^ We also considered the presence of anti-IFN-α_2_ autoantibodies, which has been associated with severe forms of the disease, as a driver of PASC.^35^ However, no link to PASC status could be established herein, as only two of the six positive patients were PASC. These results are in agreement with studies that found limited contribution of such autoantibodies to PASC,^14,36,37^ while contrasting with others that identified an association between the presence of anti-IFN-I autoantibodies and respiratory-viral PASC symptoms.^15^

The link between viral reactivation and the development of PASC was also evaluated from 6 months post-infection. TTV was related to the initial severity, exhibiting higher viral loads in patients with severe COVID-19 compared to those with mild forms, but not to the PASC status. We found no link between CMV or EBV reactivation, and disease severity or PASC status. While a previous study reported similar results regarding the absence of EBV viremia,^38^ some studies found an increase in the level of antibodies targeting EBV surface-antigens, which could indicate a recent EBV reactivation in patients with PASC.^14,25,26^ Nevertheless, the timing post-infection has to be considered, since the absence of EBV viremia at 6 months may be due to viral clearance; Su *et al.* reported a 14% viremia at diagnosis but barely detectable levels 2 to 3 months post-symptom onset^15^. Overall, our results suggest that PASC is not associated with an ongoing reactivation of persistent viruses such as EBV or CMV.

The persistence of SARS-CoV-2 in patients with PASC has been suggested by the detection of SARS-CoV-2 antigens in the blood or non-classical monocytes^23,24,39^ of patients with PASC. In addition, the presence of RNA encoding viral proteins was found in multiple tissue-biopsies obtained from these patients.^40,41^ However, in the present study, we have been unable to detect circulating SARS-CoV-2 nucleocapsid antigens in the plasma samples of the 458 included patients, regardless of the PASC status. This could be due to a lack of sensitivity of the current detection techniques, despite the use of digital ELISA herein. The reliable detection of these antigens in blood would necessitate the development of more sensitive detection kits that could help in the diagnostic of PASC.

We also measured hormonal, cytokine, and damage biomarkers that have been associated with the development of PASC.^14,15,27,28,30,34,42–48^ Although we found differences in the level of several of these proteins between the two cohorts, such differences were not associated with the PASC status within the cohorts.

PASC has been associated with impaired immune alterations in the lung^49,50^ or in PBMCs.^14,51–53^ PBMCs from patients with severe COVID-19, with and without PASC, were analyzed to assess their abundance and phenotype. In terms of subset distribution, patients with PASC tended to exhibit higher numbers of non-classical monocytes. Differences in monocyte subsets have also been reported in several previous studies in PBMCs^14,29,39^ and in the lung.^49,50^. Furthermore, using Diffcyt to define activation markers associated with PASC, we showed a greater frequency of CD4^+^ TEMRA expressing either CD57 or Tbet, while the frequency of cells expressing CXCR3 was lower in patients with PASC. Similarly, the frequency of CD8^+^ TEMRA cells expressing CXCR3 was lower in patients with PASC. This could reflect a more terminally differentiated phenotype in T cells of patients with PASC, which is in line with other studies that reported an increased expression of CD57.^52^

Subsequently, we monitored the CD4^+^ and CD8^+^ T cells response against SARS-CoV-2 structural proteins, and found that an increased frequency of N/M-responsive T cells was significantly associated with the PASC status. Importantly, TEMRA N/M-responsive CD4^+^ T cells exhibited a higher frequency of CD57 or Tbet expressing cells, and a lower frequency of CXCR3 and CD27 expressing cells, indicating that SARS-CoV-2 specific CD4^+^ T cells show the same phenotypic bias as the total CD4^+^ population. Another feature associated with PBMCs from patients with PASC was their significantly higher frequency of IFNγ secreting cells in the absence of peptide stimulation *ex vivo*, which is consistent with Krishna *et al.*, who reported increased IFNγ-secreting cells in patients with PASC 6 months post-infection, with a decreased as the disease resolved.^51^ It is worth noting that an increased secretion of IFNγ has been previously associated with disease severity.^54^ Moreover, a recent study has highlighted the potential role of this cytokine in the development of pulmonary sequelae.^49^

Finally, we used a random forest algorithm to identify parameters that could best discriminate patients with PASC from those without. The top 10 parameters identified could discriminate patients with an 80% accuracy. This analysis identified the number and duration of symptoms during the initial infection as crucial parameters, confirming findings from previous studies. ^19,55–57^ Furthermore, several parameters related to T cell phenotype and antiviral responses emerged as key factors distinguishing patients with and without PASC, highlighting the role of T cells in the development of PASC.

Our study has several limitations. Although the cohorts were diverse in age, sex, and comorbidities, they were not specifically selected for PASC assessment and represent a broad population that experienced mild-to-severe COVID-19. PASC classification relied solely on self-reported data from interviews. The sample size also limited the ability to create robust categories of persistent symptoms for detailed analysis.

The present study highlighted that the immune alterations observed in patients with PASC cannot be considered a universal feature, since they are significantly influenced by the severity of the initial COVID-19 episode and PASC heterogeneity, which encompasses a wide range of symptoms and mechanisms. Along with published data, our results confirm the challenge of establishing reliable biomarker signatures to identify patients with PASC.

## RESOURCE AVAILABILITY STATEMENT

### Lead contact

Requests for further information and resources should be directed to and will be fulfilled by the lead contact, William Mouton.

### Material availability

This study did not generate new unique reagents.

### Data and code availability

- All data reported in this paper will be shared by the lead contact upon request.
- This paper does not report original code.
- Any additional information required to reanalyze the data reported in this paper is available from the lead contact upon request.

## ACKNOWLEDGEMENTS

We thank the staff members of the Occupational Health and Medicine Department, the associated clinical research teams of Professors Charpurlat and Vanhemns for their excellent work, the members of the clinical research and innovation department for their reactivity (Hospices Civils de Lyon); and all the individuals for their participation in this clinical study. Human biological samples and associated data were obtained from NeuroBioTec (CRB HCL, Lyon France, Biobank BB-0033-00046). We thank Shanez Haouari (DRS, HCL) for help in manuscript preparation. This study has benefited funding from the Agence Nationale de Recherches sur le Sida et les Hépatites Virales (EMERGEN - ANRS - AAP Covid long 2021, numero ECTZ199036) and a partial support from the Center of Excellence in Respiratory Pathogens (CERP).

## AUTOR CONTRIBUTIONS

TW, JM, and STA participated in conceptualization,

WM, SD, MV, and SB participated in data curation,

OA and CC participated in formal analysis,

PV, TW, JM, and STA contributed to funding acquisition,

WM, SD, MV, PV, JM, TW, and STA participated in investigation

WM, SD, MV, OA, and CC contributed in methodology

JM, TW, and STA participated in project administration and supervision,

WM, SD, OA, and CC participated in visualization,

WM and SD contributed in writing the original draft,

All authors participated in reviewing & editing the final draft.

## DECLARATION OF INTERESTS

The authors declare no competing interests. This includes employment, consultancies, honoraria, stock ownership or options, expert testimony or patents received or pending, or royalties.

## SUPPLEMENTAL INFORMATION

Figures S1-S7 and Table S1-S4.

## REFERENCES

1. Lopez-Leon, S., Wegman-Ostrosky, T., Perelman, C., Sepulveda, R., Rebolledo, P.A., Cuapio, A., and Villapol, S. (2021). More than 50 long-term effects of COVID-19: a systematic review and meta-analysis. Sci. Rep. 11, 16144. 10.1038/s41598-021-95565-8.

2. Del Rio, C., Collins, L.F., and Malani, P. (2020). Long-term Health Consequences of COVID-19. JAMA 324, 1723. 10.1001/jama.2020.19719.

3. APCOVID-19 : étude nationale sur la prévalence et l’impact de l’affection post-COVID-19 https://www.santepubliquefrance.fr/etudes-et-enquetes/apcovid-19-etude-nationale-sur-la-prevalence-et-l-impact-de-l-affection-post-covid-19.

4. Aiyegbusi, O.L., Hughes, S.E., Turner, G., Rivera, S.C., McMullan, C., Chandan, J.S., Haroon, S., Price, G., Davies, E.H., Nirantharakumar, K., et al. (2021). Symptoms, complications and management of long COVID: a review. J. R. Soc. Med. 114, 428–442. 10.1177/01410768211032850.

5. Al-Aly, Z., Xie, Y., and Bowe, B. (2021). High-dimensional characterization of post-acute sequelae of COVID-19. Nature 594, 259–264. 10.1038/s41586-021-03553-9.

6. Davis, H.E., Assaf, G.S., McCorkell, L., Wei, H., Low, R.J., Re’em, Y., Redfield, S., Austin, J.P., and Akrami, A. (2021). Characterizing long COVID in an international cohort: 7 months of symptoms and their impact. EClinicalMedicine 38, 101019. 10.1016/j.eclinm.2021.101019.

7. Reese, J.T., Blau, H., Casiraghi, E., Bergquist, T., Loomba, J.J., Callahan, T.J., Laraway, B., Antonescu, C., Coleman, B., Gargano, M., et al. (2023). Generalisable long COVID subtypes: findings from the NIH N3C and RECOVER programmes. EBioMedicine 87, 104413. 10.1016/j.ebiom.2022.104413.

8. O’Mahoney, L.L., Routen, A., Gillies, C., Ekezie, W., Welford, A., Zhang, A., Karamchandani, U., Simms-Williams, N., Cassambai, S., Ardavani, A., et al. (2023). The prevalence and long-term health effects of Long Covid among hospitalised and non-hospitalised populations: A systematic review and meta-analysis. EClinicalMedicine 55, 101762. 10.1016/j.eclinm.2022.101762.

9. Subramanian, A., Nirantharakumar, K., Hughes, S., Myles, P., Williams, T., Gokhale, K.M., Taverner, T., Chandan, J.S., Brown, K., Simms-Williams, N., et al. (2022). Symptoms and risk factors for long COVID in non-hospitalized adults. Nat. Med. 28, 1706–1714. 10.1038/s41591-022-01909-w.

10. Davis, H.E., McCorkell, L., Vogel, J.M., and Topol, E.J. (2023). Long COVID: major findings, mechanisms and recommendations. Nat. Rev. Microbiol. 21, 133–146. 10.1038/s41579-022-00846-2.

11. The Lancet, null (2021). Understanding long COVID: a modern medical challenge. Lancet Lond. Engl. 398, 725. 10.1016/S0140-6736(21)01900-0.

12. Greenhalgh, T., Sivan, M., Perlowski, A., and Nikolich, J.Ž. (2024). Long COVID: a clinical update. Lancet Lond. Engl. 404, 707–724. 10.1016/S0140-6736(24)01136-X.

13. Katz, G.M., Bach, K., Bobos, P., Cheung, A., Décary, S., Goulding, S., Herridge, M.S., McNaughton, C.D., Palmer, K.S., Razak, F.A., et al. (2023). Understanding How Post-COVID-19 Condition Affects Adults and Health Care Systems. JAMA Health Forum 4, e231933. 10.1001/jamahealthforum.2023.1933.

14. Klein, J., Wood, J., Jaycox, J.R., Dhodapkar, R.M., Lu, P., Gehlhausen, J.R., Tabachnikova, A., Greene, K., Tabacof, L., Malik, A.A., et al. (2023). Distinguishing features of long COVID identified through immune profiling. Nature 623, 139–148. 10.1038/s41586-023-06651-y.

15. Su, Y., Yuan, D., Chen, D.G., Ng, R.H., Wang, K., Choi, J., Li, S., Hong, S., Zhang, R., Xie, J., et al. (2022). Multiple early factors anticipate post-acute COVID-19 sequelae. Cell 185, 881–895.e20. 10.1016/j.cell.2022.01.014.

16. Parotto, M., Gyöngyösi, M., Howe, K., Myatra, S.N., Ranzani, O., Shankar-Hari, M., and Herridge, M.S. (2023). Post-acute sequelae of COVID-19: understanding and addressing the burden of multisystem manifestations. Lancet Respir. Med. 11, 739–754. 10.1016/S2213-2600(23)00239-4.

17. Saadatian-Elahi, M., Picot, V., Hénaff, L., Pradel, F.K., Escuret, V., Dananché, C., Elias, C., Endtz, H.P., and Vanhems, P. (2020). Protocol for a prospective, observational, hospital-based multicentre study of nosocomial SARS-CoV-2 transmission: NOSO-COR Project. BMJ Open 10, e039088. 10.1136/bmjopen-2020-039088.

18. Trouillet-Assant, S., Albert Vega, C., Bal, A., Nazare, J.A., Fascia, P., Paul, A., Massardier-Pilonchery, A., D Aubarede, C., Guibert, N., Pitiot, V., et al. (2020). Assessment of serological techniques for screening patients for COVID-19 (COVID-SER): a prospective, multicentric study. BMJ Open 10, e041268. 10.1136/bmjopen-2020-041268.

19. Khanafer, N., Henaff, L., Bennia, S., Termoz, A., Chapurlat, R., Escuret, V., Proriol, M., Duvert, F., Mena, C., Planckaert, C., et al. (2023). Factors Associated with Long COVID-19 in a French Multicentric Prospective Cohort Study. Int. J. Environ. Res. Public. Health 20, 6678. 10.3390/ijerph20176678.

20. Mohandas, S., Jagannathan, P., Henrich, T.J., Sherif, Z.A., Bime, C., Quinlan, E., Portman, M.A., Gennaro, M., Rehman, J., and RECOVER Mechanistic Pathways Task Force (2023). Immune mechanisms underlying COVID-19 pathology and post-acute sequelae of SARS-CoV-2 infection (PASC). eLife 12, e86014. 10.7554/eLife.86014.

21. Binswanger, I.A., Palmer-Toy, D.E., Barrow, J.C., Narwaney, K.J., Bruxvoort, K.J., Kraus, C.R., Lyons, J.A., Lam, J.A., and Glanz, J.M. (2024). Assessing the association between antibody status and symptoms of long COVID: A multisite study. PloS One 19, e0304262. 10.1371/journal.pone.0304262.

22. Jansen, E.B., Ostadgavahi, A.T., Hewins, B., Buchanan, R., Thivierge, B.M., Sganzerla Martinez, G., Goncin, U., Francis, M.E., Swan, C.L., Scruten, E., et al. (2024). PASC (Post Acute Sequelae of COVID-19) is associated with decreased neutralizing antibody titers in both biological sexes and increased ANG-2 and GM-CSF in females. Sci. Rep. 14, 9854. 10.1038/s41598-024-60089-4.

23. Swank, Z., Senussi, Y., Manickas-Hill, Z., Yu, X.G., Li, J.Z., Alter, G., and Walt, D.R. (2023). Persistent Circulating Severe Acute Respiratory Syndrome Coronavirus 2 Spike Is Associated With Post-acute Coronavirus Disease 2019 Sequelae. Clin. Infect. Dis. Off. Publ. Infect. Dis. Soc. Am. 76, e487–e490. 10.1093/cid/ciac722.

24. Swank, Z., Borberg, E., Chen, Y., Senussi, Y., Chalise, S., Manickas-Hill, Z., Yu, X.G., Li, J.Z., Alter, G., Henrich, T.J., et al. (2024). Measurement of circulating viral antigens post-SARS-CoV-2 infection in a multicohort study. Clin. Microbiol. Infect. Off. Publ. Eur. Soc. Clin. Microbiol. Infect. Dis., S1198-743X(24)00432-4. 10.1016/j.cmi.2024.09.001.

25. Gold, J.E., Okyay, R.A., Licht, W.E., and Hurley, D.J. (2021). Investigation of Long COVID Prevalence and Its Relationship to Epstein-Barr Virus Reactivation. Pathog. Basel Switz. 10, 763. 10.3390/pathogens10060763.

26. Peluso, M.J., Deveau, T.-M., Munter, S.E., Ryder, D., Buck, A., Beck-Engeser, G., Chan, F., Lu, S., Goldberg, S.A., Hoh, R., et al. (2023). Chronic viral coinfections differentially affect the likelihood of developing long COVID. J. Clin. Invest. 133, e163669. 10.1172/JCI163669.

27. Schultheiß, C., Willscher, E., Paschold, L., Gottschick, C., Klee, B., Henkes, S.-S., Bosurgi, L., Dutzmann, J., Sedding, D., Frese, T., et al. (2022). The IL-1β, IL-6, and TNF cytokine triad is associated with post-acute sequelae of COVID-19. Cell Rep. Med. 3, 100663. 10.1016/j.xcrm.2022.100663.

28. Peluso, M.J., Lu, S., Tang, A.F., Durstenfeld, M.S., Ho, H.-E., Goldberg, S.A., Forman, C.A., Munter, S.E., Hoh, R., Tai, V., et al. (2021). Markers of Immune Activation and Inflammation in Individuals With Postacute Sequelae of Severe Acute Respiratory Syndrome Coronavirus 2 Infection. J. Infect. Dis. 224, 1839–1848. 10.1093/infdis/jiab490.

29. Phetsouphanh, C., Darley, D.R., Wilson, D.B., Howe, A., Munier, C.M.L., Patel, S.K., Juno, J.A., Burrell, L.M., Kent, S.J., Dore, G.J., et al. (2022). Immunological dysfunction persists for 8 months following initial mild-to-moderate SARS-CoV-2 infection. Nat. Immunol. 23, 210–216. 10.1038/s41590-021-01113-x.

30. Fleischer, M., Szepanowski, F., Mausberg, A.K., Asan, L., Uslar, E., Zwanziger, D., Volbracht, L., Stettner, M., and Kleinschnitz, C. (2024). Cytokines (IL1β, IL6, TNFα) and serum cortisol levels may not constitute reliable biomarkers to identify individuals with post-acute sequelae of COVID-19. Ther. Adv. Neurol. Disord. 17, 17562864241229567. 10.1177/17562864241229567.

31. Pink, I., Hennigs, J.K., Ruhl, L., Sauer, A., Boblitz, L., Huwe, M., Fuge, J., Falk, C.S., Pietschmann, T., de Zwaan, M., et al. (2024). Blood T cell phenotypes correlate with fatigue severity in post-acute sequelae of COVID-19. Infection 52, 513–524. 10.1007/s15010-023-02114-8.

32. Woodrow, M., Carey, C., Ziauddeen, N., Thomas, R., Akrami, A., Lutje, V., Greenwood, D.C., and Alwan, N.A. (2023). Systematic Review of the Prevalence of Long COVID. Open Forum Infect. Dis. 10, ofad233. 10.1093/ofid/ofad233.

33. Al-Aly, Z., and Topol, E. (2024). Solving the puzzle of Long Covid. Science 383, 830–832. 10.1126/science.adl0867.

34. Peluso, M.J., Sans, H.M., Forman, C.A., Nylander, A.N., Ho, H.-E., Lu, S., Goldberg, S.A., Hoh, R., Tai, V., Munter, S.E., et al. (2022). Plasma Markers of Neurologic Injury and Inflammation in People With Self-Reported Neurologic Postacute Sequelae of SARS-CoV-2 Infection. Neurol. Neuroimmunol. Neuroinflammation 9, e200003. 10.1212/NXI.0000000000200003.

35. Goncalves, D., Mezidi, M., Bastard, P., Perret, M., Saker, K., Fabien, N., Pescarmona, R., Lombard, C., Walzer, T., Casanova, J.-L., et al. (2021). Antibodies against type I interferon: detection and association with severe clinical outcome in COVID-19 patients. Clin. Transl. Immunol. 10, e1327. 10.1002/cti2.1327.

36. Peluso, M.J., Mitchell, A., Wang, C.Y., Takahashi, S., Hoh, R., Tai, V., Durstenfeld, M.S., Hsue, P.Y., Kelly, J.D., Martin, J.N., et al. (2023). Low Prevalence of Interferon α Autoantibodies in People Experiencing Symptoms of Post-Coronavirus Disease 2019 (COVID-19) Conditions, or Long COVID. J. Infect. Dis. 227, 246–250. 10.1093/infdis/jiac372.

37. Achleitner, M., Mair, N.K., Dänhardt, J., Kardashi, R., Puhan, M.A., Abela, I.A., Toepfner, N., de With, K., Kanczkowski, W., Jarzebska, N., et al. (2024). Absence of Type I Interferon Autoantibodies or Significant Interferon Signature Alterations in Adults With Post-COVID-19 Syndrome. Open Forum Infect. Dis. 11, ofad641. 10.1093/ofid/ofad641.

38. Hoeggerl, A.D., Nunhofer, V., Lauth, W., Badstuber, N., Held, N., Zimmermann, G., Grabmer, C., Weidner, L., Jungbauer, C., Lindlbauer, N., et al. (2023). Epstein-Barr virus reactivation is not causative for post-COVID-19-syndrome in individuals with asymptomatic or mild SARS-CoV-2 disease course. BMC Infect. Dis. 23, 800. 10.1186/s12879-023-08820-w.

39. Patterson, B.K., Francisco, E.B., Yogendra, R., Long, E., Pise, A., Rodrigues, H., Hall, E., Herrera, M., Parikh, P., Guevara-Coto, J., et al. (2021). Persistence of SARS CoV-2 S1 Protein in CD16+ Monocytes in Post-Acute Sequelae of COVID-19 (PASC) up to 15 Months Post-Infection. Front. Immunol. 12, 746021. 10.3389/fimmu.2021.746021.

40. Peluso, M.J., Ryder, D., Flavell, R.R., Wang, Y., Levi, J., LaFranchi, B.H., Deveau, T.-M., Buck, A.M., Munter, S.E., Asare, K.A., et al. (2024). Tissue-based T cell activation and viral RNA persist for up to 2 years after SARS-CoV-2 infection. Sci. Transl. Med. 16, eadk3295. 10.1126/scitranslmed.adk3295.

41. Zuo, W., He, D., Liang, C., Du, S., Hua, Z., Nie, Q., Zhou, X., Yang, M., Tan, H., Xu, J., et al. (2024). The persistence of SARS-CoV-2 in tissues and its association with long COVID symptoms: a cross-sectional cohort study in China. Lancet Infect. Dis. 24, 845–855. 10.1016/S1473-3099(24)00171-3.

42. Bark, L., Larsson, I.-M., Wallin, E., Simrén, J., Zetterberg, H., Lipcsey, M., Frithiof, R., Rostami, E., and Hultström, M. (2023). Central nervous system biomarkers GFAp and NfL associate with post-acute cognitive impairment and fatigue following critical COVID-19. Sci. Rep. 13, 13144. 10.1038/s41598-023-39698-y.

43. Alaedini, A., Lightman, S., and Wormser, G.P. (2024). Is Low Cortisol a Marker of Long COVID? Am. J. Med. 137, 564–565. 10.1016/j.amjmed.2024.03.013.

44. Lai, Y.-J., Liu, S.-H., Manachevakul, S., Lee, T.-A., Kuo, C.-T., and Bello, D. (2023). Biomarkers in long COVID-19: A systematic review. Front. Med. 10, 1085988. 10.3389/fmed.2023.1085988.

45. Williams, E.S., Martins, T.B., Shah, K.S., Hill, H.R., Coiras, M., Spivak, A.M., and Planelles, V. (2022). Cytokine Deficiencies in Patients with Long-COVID. J. Clin. Cell. Immunol. 13, 672.

46. 46. Queiroz, M.A.F., Neves, P.F.M. das, Lima, S.S., Lopes, J. da C., Torres, M.K. da S., Vallinoto, I.M.V.C., Bichara, C.D.A., Dos Santos, E.F., de Brito, M.T.F.M., da Silva, A.L.S., et al. (2022). Cytokine Profiles Associated With Acute COVID-19 and Long COVID-19 Syndrome. Front. Cell. Infect. Microbiol. 12, 922422. 10.3389/fcimb.2022.922422.

47. Hanson, B.A., Visvabharathy, L., Ali, S.T., Kang, A.K., Patel, T.R., Clark, J.R., Lim, P.H., Orban, Z.S., Hwang, S.S., Mattoon, D., et al. (2022). Plasma Biomarkers of Neuropathogenesis in Hospitalized Patients With COVID-19 and Those With Postacute Sequelae of SARS-CoV-2 Infection. Neurol. Neuroimmunol. Neuroinflammation 9, e1151. 10.1212/NXI.0000000000001151.

48. Spanos, M., Shachar, S., Sweeney, T., Lehmann, H.I., Gokulnath, P., Li, G., Sigal, G.B., Nagaraj, R., Bathala, P., Rana, F., et al. (2022). Elevation of neural injury markers in patients with neurologic sequelae after hospitalization for SARS-CoV-2 infection. iScience 25, 104833. 10.1016/j.isci.2022.104833.

49. 49. Li, C., Qian, W., Wei, X., Narasimhan, H., Wu, Y., Arish, M., Cheon, I.S., Tang, J., de Almeida Santos, G., Li, Y., et al. (2024). Comparative single-cell analysis reveals IFN-γ as a driver of respiratory sequelae after acute COVID-19. Sci. Transl. Med. 16, eadn0136. 10.1126/scitranslmed.adn0136.

50. Vijayakumar, B., Boustani, K., Ogger, P.P., Papadaki, A., Tonkin, J., Orton, C.M., Ghai, P., Suveizdyte, K., Hewitt, R.J., Desai, S.R., et al. (2022). Immuno-proteomic profiling reveals aberrant immune cell regulation in the airways of individuals with ongoing post-COVID-19 respiratory disease. Immunity 55, 542–556.e5. 10.1016/j.immuni.2022.01.017.

51. Krishna, B.A., Lim, E.Y., Metaxaki, M., Jackson, S., Mactavous, L., NIHR BioResource, Lyons, P.A., Doffinger, R., Bradley, J.R., Smith, K.G.C., et al. (2024). Spontaneous, persistent, T cell-dependent IFN-γ release in patients who progress to Long Covid. Sci. Adv. 10, eadi9379. 10.1126/sciadv.adi9379.

52. Galán, M., Vigón, L., Fuertes, D., Murciano-Antón, M.A., Casado-Fernández, G., Domínguez-Mateos, S., Mateos, E., Ramos-Martín, F., Planelles, V., Torres, M., et al. (2022). Persistent Overactive Cytotoxic Immune Response in a Spanish Cohort of Individuals With Long-COVID: Identification of Diagnostic Biomarkers. Front. Immunol. 13, 848886. 10.3389/fimmu.2022.848886.

53. Rodriguez, L., Tan, Z., Lakshmikanth, T., Wang, J., Barcenilla, H., Swank, Z., Zuo, F., Abolhassani, H., Pavlovitch-Bedzyk, A.J., Wang, C., et al. (2024). Restrained memory CD8^+^ T cell responses favors viral persistence and elevated IgG responses in patients with severe Long COVID. Preprint at Infectious Diseases (except HIV/AIDS), 10.1101/2024.02.11.24302636 https://doi.org/10.1101/2024.02.11.24302636.

54. Peluso, M.J., Deitchman, A.N., Torres, L., Iyer, N.S., Munter, S.E., Nixon, C.C., Donatelli, J., Thanh, C., Takahashi, S., Hakim, J., et al. (2021). Long-term SARS-CoV-2-specific immune and inflammatory responses in individuals recovering from COVID-19 with and without post-acute symptoms. Cell Rep. 36, 109518. 10.1016/j.celrep.2021.109518.

55. Sudre, C.H., Murray, B., Varsavsky, T., Graham, M.S., Penfold, R.S., Bowyer, R.C., Pujol, J.C., Klaser, K., Antonelli, M., Canas, L.S., et al. (2021). Attributes and predictors of long COVID. Nat. Med. 27, 626–631. 10.1038/s41591-021-01292-y.

56. Wu, Y., Sawano, M., Wu, Y., Shah, R.M., Bishop, P., Iwasaki, A., and Krumholz, H.M. (2024). Factors Associated With Long COVID: Insights From Two Nationwide Surveys. Am. J. Med. 137, 515–519. 10.1016/j.amjmed.2024.02.032.

57. Tsampasian, V., Elghazaly, H., Chattopadhyay, R., Debski, M., Naing, T.K.P., Garg, P., Clark, A., Ntatsaki, E., and Vassiliou, V.S. (2023). Risk Factors Associated With Post-COVID-19 Condition: A Systematic Review and Meta-analysis. JAMA Intern. Med. 183, 566–580. 10.1001/jamainternmed.2023.0750.

58. Bastard, P., Michailidis, E., Hoffmann, H.-H., Chbihi, M., Le Voyer, T., Rosain, J., Philippot, Q., Seeleuthner, Y., Gervais, A., Materna, M., et al. (2021). Auto-antibodies to type I IFNs can underlie adverse reactions to yellow fever live attenuated vaccine. J. Exp. Med. 218, e20202486. 10.1084/jem.20202486.

59. Mouton, W., Compagnon, C., Saker, K., Daniel, S., Djebali, S., Lacoux, X., Pozzetto, B., Oriol, G., Laubreton, D., Prieux, M., et al. (2021). Specific detection of memory T-cells in COVID-19 patients using standardized whole-blood Interferon gammarelease assay. Eur. J. Immunol. 51, 3239– 3242. 10.1002/eji.202149296.

60. diffcyt Bioconductor. http://bioconductor.org/packages/diffcyt/.

61. Weber, L.M., Nowicka, M., Soneson, C., and Robinson, M.D. (2019). diffcyt: Differential discovery in high-dimensional cytometry via high-resolution clustering. Commun. Biol. 2, 183. 10.1038/s42003-019-0415-5.

